# Variable Genome Evolution in Fungi After Transposon-Mediated Amplification of a Housekeeping Gene

**DOI:** 10.1101/550798

**Authors:** Braham Dhillon, Gert H. J. Kema, Richard Hamelin, Burt H. Bluhm, Stephen B. Goodwin

## Abstract

**Background:** Transposable elements (TEs) can be key drivers of evolution, but the mechanisms and scope of how they impact gene and genome function are largely unknown. Previous analyses revealed that TE-mediated gene amplifications can have variable effects on fungal genomes, from inactivation of function to production of multiple active copies. For example, a DNA methyltransferase gene in the wheat pathogen *Zymoseptoria tritici* (synonym *Mycosphaerella graminicola*) was amplified to tens of copies, all of which were inactivated by Repeat-Induced Point mutation (RIP) including the original, resulting in loss of cytosine methylation. In another wheat pathogen, *Pyrenophora tritici-repentis*, a histone H3 gene was amplified to tens of copies with little evidence of RIP, leading to many potentially active copies. To further test the effects of transposon-aided gene amplifications on genome evolution and architecture, the repetitive fraction of the significantly expanded *Pseudocercospora fijiensis* genome was analyzed in greater detail.

**Results:** These analyses identified a housekeeping gene, histone H3, which was captured and amplified to hundreds of copies by a *hAT* DNA transposon, all of which were inactivated by RIP, except for the original. In *P. fijiensis* the original H3 gene probably was not protected from RIP, but most likely was maintained intact due to strong purifying selection. Comparative analyses revealed that a similar event occurred in five additional genomes representing the fungal genera *Cercospora, Pseudocercospora* and *Sphaerulina*.

**Conclusions:** These results indicate that the interplay of TEs and RIP can result in different and unpredictable fates of amplified genes, with variable effects on gene and genome evolution.

## Background

Transposable elements (TEs) or mobile genetic elements are nucleic acid entities that can excise and insert randomly in a genome. TEs have been detected in the genomes of all prokaryotic and eukaryotic organisms examined so far [1], and have been rightly labeled as ‘drivers of genome evolution’ [2] due to their direct and indirect impacts on genes and genomes. Several lines of evidence point to their pivotal role in important processes across the tree of life. For example, *Alu* elements, at more than a million copies comprising 11% of the human genome, are a major contributor to primate genome evolution and the standing genetic diversity in human populations [3]. Some major effects of *Alu*-element amplifications include alterations of gene expression from insertions near gene promotors, insertional mutagenesis and repeat-mediated non-homologous recombination that can lead to disease, and ‘exonization’ of *Alu* elements yielding alternative splicing of transcripts. All of these likely have played an important role in the evolution of humans and other primates [3].

TEs are major contributors to genetic diversity in populations and have been linked to major phenotypic changes in plant morphology. Natural and artificial selection can act on this variation to favor new morphotypes. For example, the transition in plant architecture from highly branched teosinte, the ancestor of modern corn, to apically dominant corn is controlled in part by a quantitative trait locus (QTL), *tb1* [4], whose expression is modulated by *Hopscotch*, a retrotransposon enhancer located 65 kb upstream of the gene [5]. This change in morphology predates the start of agriculture (10,000 years ago) and provided early agriculturalists with existing variation that could be selected from within populations. Similarly, a change in tomato fruit shape from round to elongate was initiated by a retrotransposon-mediated gene duplication of the SUN locus. This rearrangement introduced new upstream cis elements, which increased the expression of SUN, thereby causing the change in morphology [6].

In fungal and oomycete plant pathogens, besides modulating genome size (e.g., *Blumeria graminis* [7], *Pseudocercospora fijiensis*; [8]) and genome architecture (*Leptosphaeria maculans*; [9]), TEs also can be associated with plant-pathogen interactions through modulation of effectors and secondary metabolite gene clusters. TE-rich genomic islands in expanded fungal (*P. fijiensis, L. maculans*) and oomycete (*Phytophthora infestans*) genomes carry genes that code for lineage-specific, putative small, secreted proteins. In the barley powdery mildew pathogen, *B. graminis* f. sp. *hordei*, the amplification and diversification of an avirulence gene, AVRk1 has been attributed to a Long Interspersed Nuclear Element (LINE) TE [10]. Recently, it was shown that the sequence for this gene family of avirulence effectors was derived from the LINE TE [11]. Other fungal genome components, such as telomeres in *Magnaporthe oryzae* AVR-Pita [12] and lineage-specific chromosomes in *Fusarium oxysporum* [13] also are enriched in pathogenicity factors and TEs. Recently, TEs have been shown to be implicated in gain and/or loss of host-specific effector genes in *M. oryzae* [14].

Universal mechanisms exist that can minimize the deleterious impacts of TEs on host genomes. Post-transcriptional silencing and DNA methylation are two primary methods that limit the activity of TEs in genomes. The genetic network employed to silence the TEs also can be context dependent. In germline cells, piRNA, a specific class of small, non-coding RNA, is responsible for the epigenetic and post-transcriptional silencing of TEs [15]. Other genome-defense mechanisms are unique to specific organisms, e.g., Repeat-Induced Point mutation or RIP [16] has been described only in fungi. The RIP machinery can recognize repetitive sequences, that are approximately 400 bp or longer with identity of 80% or more, and introduce random transition (cytosine to thymine) mutations during each meiotic cycle in *Neurospora crassa* [17] and many other fungi. These mutations generate premature stop codons within TE-encoded genes that prevent translation of the proteins required for movement, thus rendering the transposon immobile. This abundance of transition mutations also skews the GC content of the sequence and makes it possible to identify signatures of RIP *in silico* [18].

A side effect of RIP is that the required machinery does not discriminate between functional genes and TEs; any sequence in a genome that is repetitive can be targeted, which occasionally causes unexpected effects. For example, a single-copy DNA methyltransferase (DNMT) gene in the wheat pathogen *Zymoseptoria tritici* (previously known as *Mycosphaerella graminicola*) was amplified to 23 copies and became a target for RIP. All of the DNMT sequences in the genome, including the original copy, were inactivated by RIP-introduced transition mutations [19]. A genome-wide assay for cytosine methylation revealed that it was lacking in the *Z. tritici* genome [19], but present in close relatives that possessed an intact copy of the DNMT gene. Those species are thought to have diverged from *Z. tritici* within the past 10,000 years [20], hence this change appears to be very recent. In another wheat pathogen, *Pyrenophora tritici-repentis* (Ptr), the histone H3 gene was captured as part of a *hAT* DNA transposon and amplified to 26 copies in the genome [21]. However, in contrast to *Z. tritici*, 23 of the histone H3 copies in the Ptr genome appeared to code for a functional protein [21] yielding multiple active copies of the gene. These two fungi are in different taxonomic orders of the class Dothideomycetes, and demonstrate that the fates of repeated sequences can vary, with different and unpredictable effects on gene and genome evolution.

The two examples described above define the extremes of the possible outcomes of TE-mediated gene amplification events in fungi with RIP, where either all the copies, including the original, can be inactivated leading to loss of gene function, or none of the copies being affected by RIP leading to multiple functional genes. To test whether similar gene amplifications are common in the Dothideomycetes, we conducted a genome-wide search in multiple sequenced species to quantify the prevalence of such events, and to investigate whether TE-associated gene amplifications occur commonly with large effects on gene and genome evolution. These analyses identified an amplification event in a fungal clade (*Cercospora* / *Pseudocercospora* / *Sphaerulina*) that fits between the spectrum of events bounded by the two extremes described above. In this newly described amplification event, the original gene was maintained, presumably due to selection, whereas all the amplified copies were targeted and inactivated by RIP, thus yielding very different outcomes from three similar gene amplifications in fungi.

## Results

### Repeats carrying histone H3-like sequence occur exclusively in AT-rich blocks

Analysis of the repetitive fraction of the *P. fijiensis* expanded genome revealed an abundance of histone H3-like sequences. A similarity search using the histone H3 protein sequence revealed a total of 784 H3-like copies that were found exclusively in repetitive regions across 28 scaffolds, with the original histone H3 gene located on scaffold 6 (Figure 1). All H3-like sequences were identified in one repeat family, with a total of 1,579 members, many of which were incomplete but overlapping and could be merged into 920 contigs, of which 471 copies contained H3-like sequences that accounted for 4.1% (3 Mb) of the *P. fijiensis* genome. Each repetitive element contained one or two copies of the H3-like sequences giving 784 copies in total. All these repeat elements were compartmentalized in the AT-rich blocks [8] that were identified previously in the *P. fijiensis* genome.

**Fig 1.**
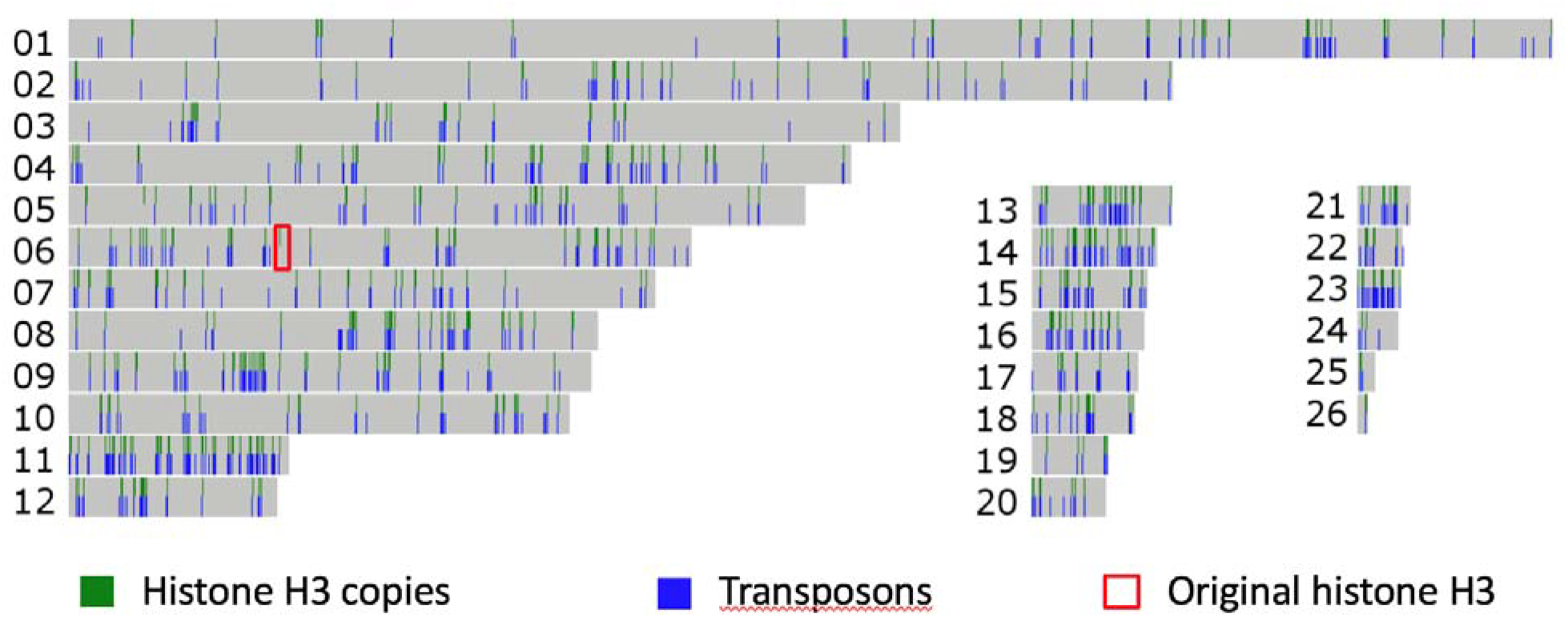
Distribution of histone H3-like sequences in the *Pseudocercospora fijiensis* genome. A total of 784 histone H3-like sequences distributed among 471 *hAT* repetitive elements along 26 scaffolds (the two smallest were excluded) in the *P. fijiensis* genome are shown. All of the H3-like sequences, except for the original, are found in repeats. For each scaffold, histone H3 copies are indicated in the top half and all transposons in the bottom. Scaffold lengths are scaled to size and the original histone H3 gene is indicated with a box

Most (90%) of the histone H3-like sequences were truncated, i.e., had lengths that ranged between 71-100 amino acids (AAs), which is 52-73% of the original histone H3 protein length of 136 AAs (Figure 2). When two histone H3-like copies were present within the same element they had lengths that differed by an insertion of 16 AAs in the second copy, which was not present in the original copy on scaffold 6.

**Fig 2.**
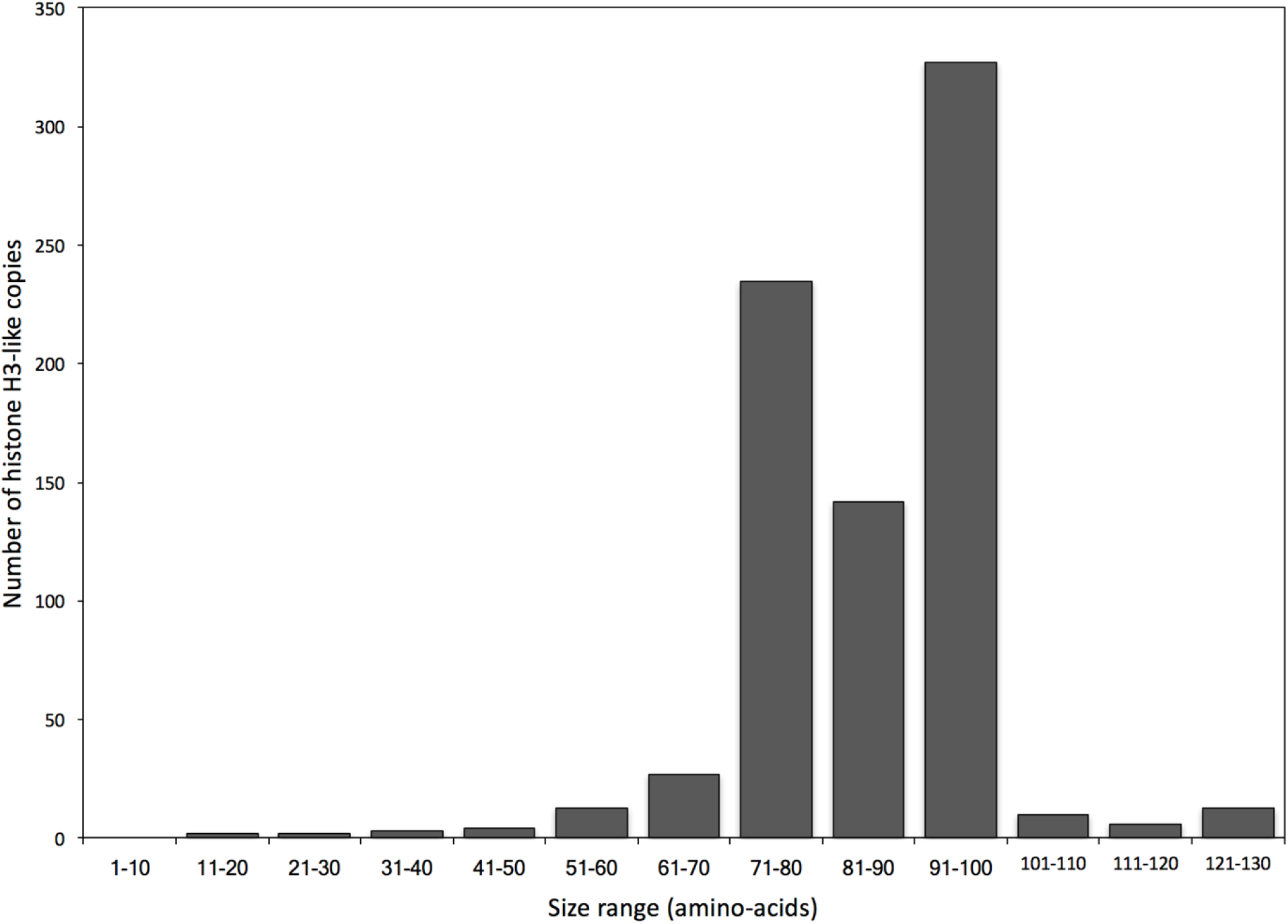
Length distribution of histone H3 gene copies in the *Pseudocercospora fijiensis* genome. Length distribution of 784 histone H3-like copies in the *P. fijiensis* genome showed that most were truncated. Only 11 (1.4%) of the sequences were near the full length (i.e., >90% of the full-length sequence), as compared to the original histone H3 protein of 136 amino acids

### A complete *hAT* DNA transposon carries the histone H3-like sequence

Annotation of the repetitive sequences flanking the histone H3-like copies identified a *hAT* transposase domain, the hallmark of *hAT* DNA transposable elements [22]. The *hAT* domain was present in 277 (59 %) of the aforementioned 471 repetitive elements that contained the histone H3-like sequence. Based on element length distribution, a group of 133 repetitive elements, with lengths ranging from 9.5 to 9.9 kb, was defined as the full-length set (Figure 3). As compared to the full-length elements, 39% of the repeat elements were considered truncated, i.e., they contained less than 50% of the full-length element (Figure 3). Both the merged repeat dataset (920 sequences) and the full-length subset (133 repeats) were used to assess the dinucleotide bias introduced by RIP. A clear CpA to TpA dinucleotide bias was observed suggesting the presence of RIP in the *P. fijiensis* genome (Figure 4). The full-length repeat set was then utilized to search for the structural features of DNA transposons, such as terminal inverted repeats (TIRs) and target site duplications (TSDs). In addition to element length and presence of *hAT* domain, the occurrence of intact TIRs and identical TSDs was used to define a repeat subset of 99 complete repetitive elements with all of these characteristics. The nucleotide composition of the 20-bp TIR sequence was well conserved across the 99-repeat set (Figure 5). Two sites in the TIR displayed the characteristic transition mutations and CpA to TpA dinucleotide bias introduced by RIP. The TSD was 8 bp in length, which is a characteristic feature of *hAT* DNA transposons [22]. No bias in insertion site was identified based on analysis of the TSD nucleotide compositions (Figure 5) of the 99 complete repetitive elements.

**Fig 3.**
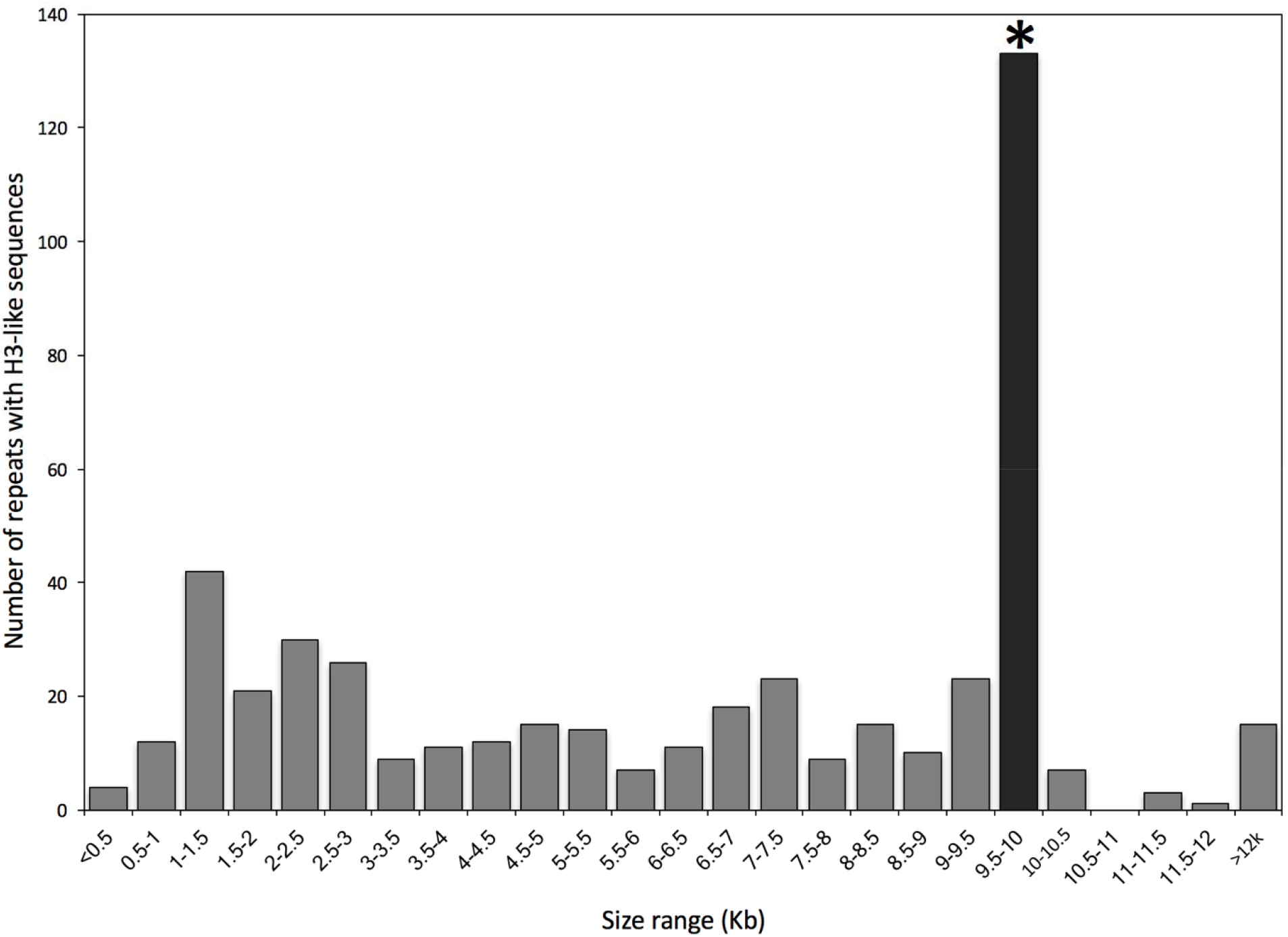
Histogram of the lengths of repetitive elements in the *Pseudocercospora fijiensis* genome that carry histone H3-like sequences. A total of 471 repetitive elements carry the H3-like sequences. Full-length repetitive elements (133) are ∼9.5 kb in length (black bar marked by an asterisk) and were used to identify the terminal inverted repeat (TIR) sequence

**Fig 4.**
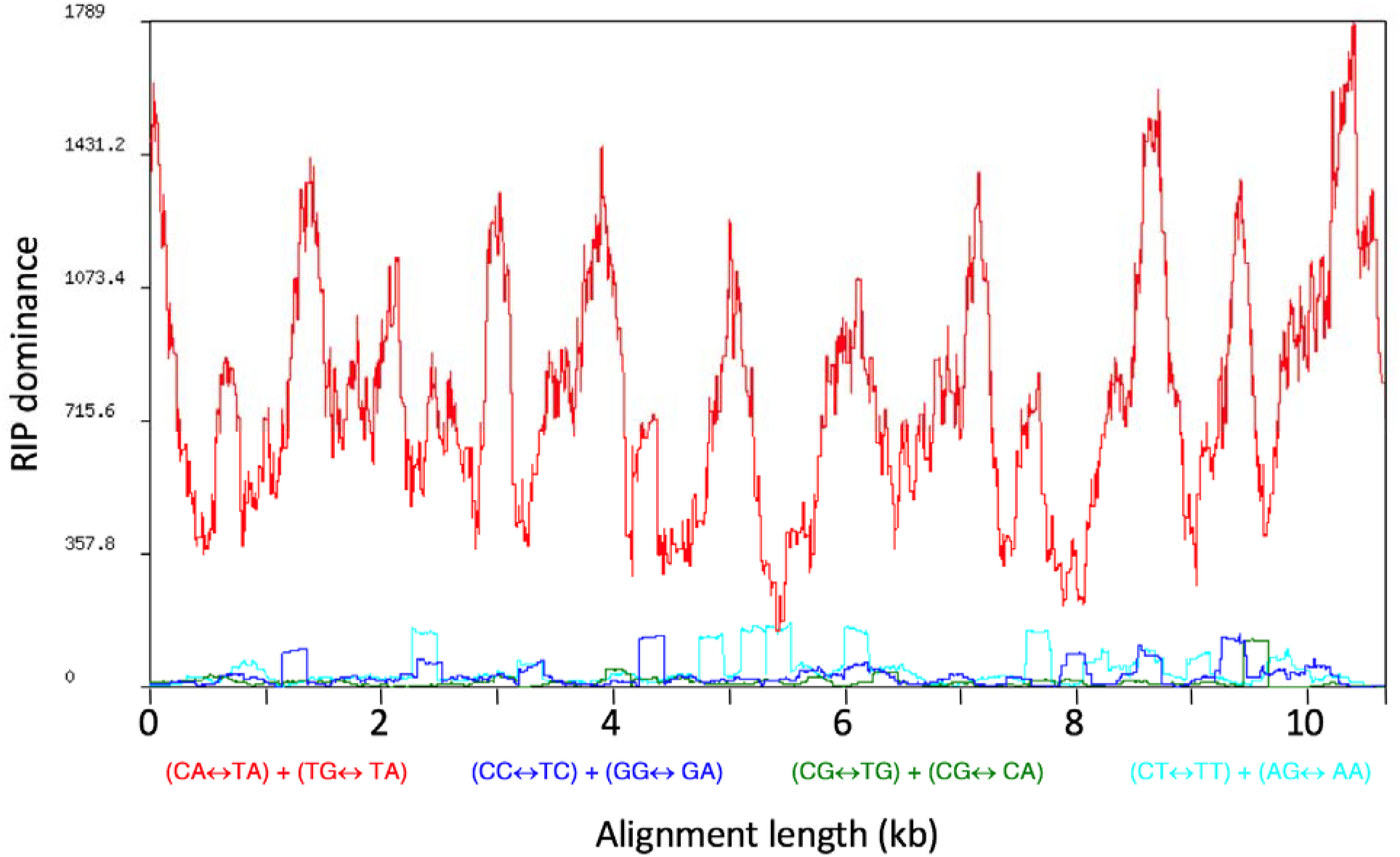
Analysis of dinucleotide bias in the *hAT* DNA transposons in the *Pseudocercospora fijiensis* genome that carry a histone H3-like sequence. Alignment of 133 bona fide *hAT* DNA transposons showed a clear CA↔TA repeat-induced point mutation (RIP) bias. RIP dominance is the ratio of one dinucleotide (CA↔TA) to the sum of the other three (CT↔TT + CG↔TG + CC↔TC)

**Fig 5.**
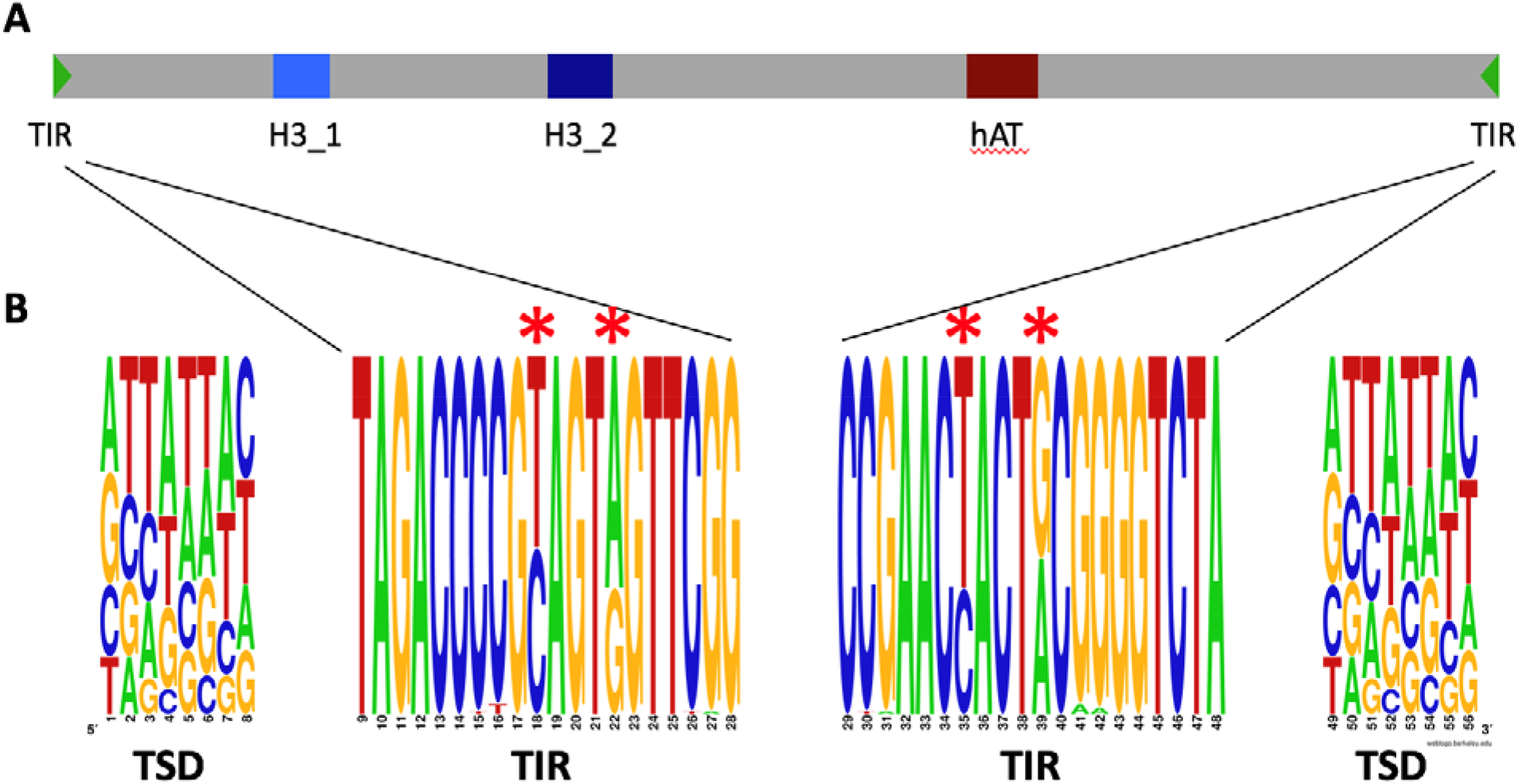
Structural organization of the *hAT* family of repetitive elements in the *Pseudocercospora fijiensis* genome that contain the histone H3-like copies. **A**) Each element had one to two copies of the H3-like sequence. **B**) A 20-bp terminal inverted repeat (TIR) at the end of each complete element was flanked by an 8-bp target site duplication (TSD) in the host DNA. These consensus sequences were derived from 99 full-length *hAT* elements that met four criteria: element length of ∼9.5 kb; presence of *hAT* domain; intact TIRs; and identical or nearly identical TSDs. Two positions within the TIRs showed the T to C and G to A transitions (indicated by an asterisk) that are characteristic of RIP.

### Transduplication of histone H3 coding sequence into a hAT DNA transposon

Co-occurrence of the histone H3-like sequence with the *hAT* DNA transposon element was investigated to test whether the complete, genomic histone gene or the transcribed coding sequence was acquired. The original genomic copy of the histone H3 gene in *P. fijiensis* has three exons (two introns), with the coding sequence (CDS) being 411 nucleotides (136 AAs) in length. Most (94%) of the H3-like copies in the genome had similarity only to the third exon of the histone H3 gene and there were few H3-like sequences spanning one (n=23, 4%) and two (n=15, 2%) exon-exon junctions, while a search using the histone HMM profile only generated partial matches to the third exon in the full-length repetitive element dataset. However, in the consensus sequence generated from *in silico* deconvolution of the RIP-introduced transition mutations, a single exon-exon junction could be recovered (Figure 6). Moreover, a near full-length histone H3 protein sequence containing the two exon-exon junctions without the intron, could be resolved when the deRIP consensus sequence was edited manually to remove all stop codons (Figure 6). The presence of histone exon-exon junctions and the absence of any intron sequence in the duplicated copies suggest that a histone H3 transcript or retrocopy, rather than a genomic copy, was captured by the *hAT* DNA transposon.

**Fig 6.**
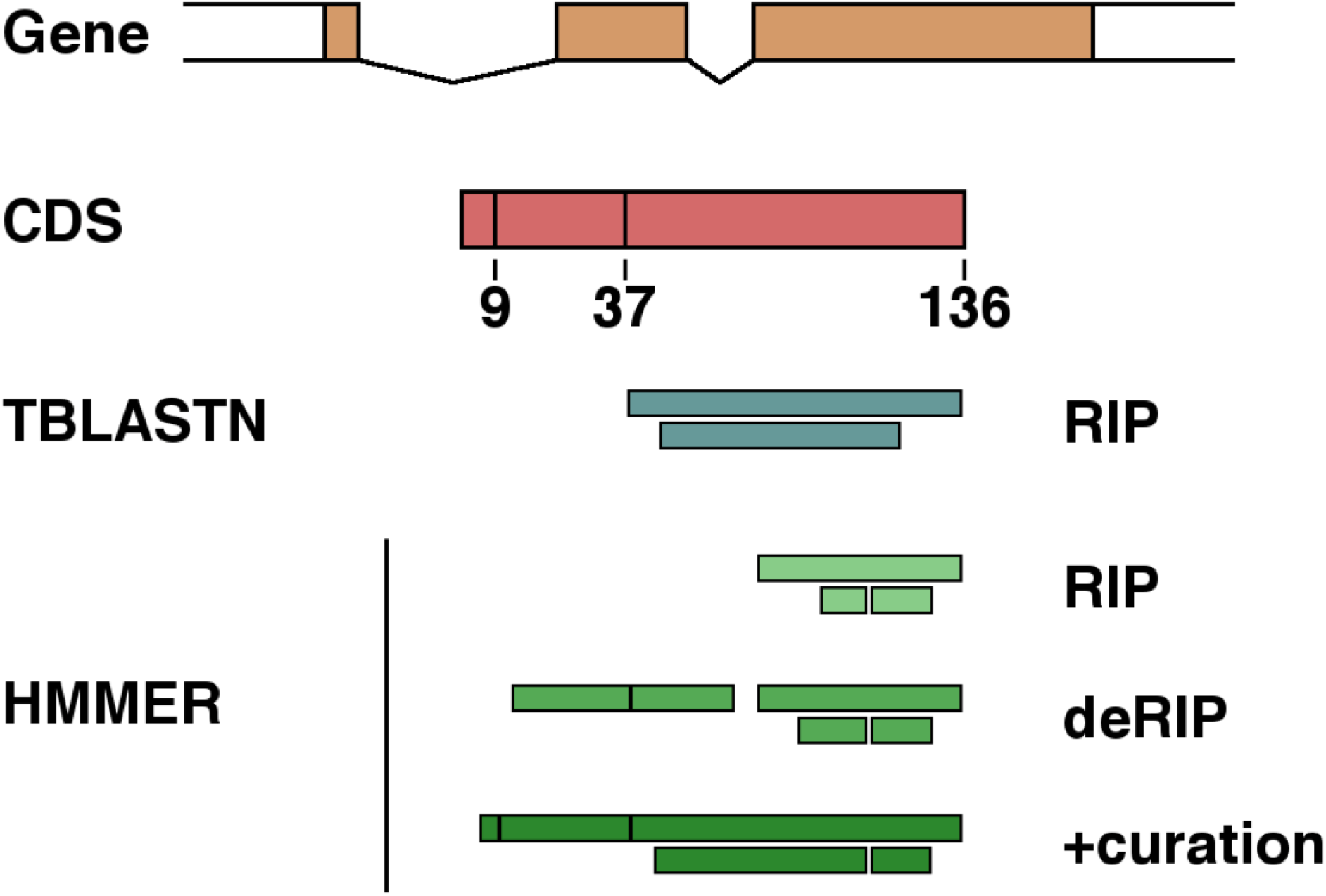
Transcribed histone H3 coding sequence was captured by an *hAT* DNA transposon. Presence of exon-exon boundaries was used as an indication of transcribed coding sequence capture by the DNA transposable elements. Initially, tBLASTn identified the histone H3 copies in the DNA transposon. The initial search was refined by using histone hmm profiles with HMMER. The original sequences had very poor H3 domain hits due to numerous changes caused by repeat-induced point mutation (RIP). The identification of H3-like sequence in deRIPped elements was substantially better. However, to obtain the best results, i.e., all exon-exon boundaries were evident, the deRIPped sequences were curated manually by removing the remaining stop codons that resulted when RIP occurred in all copies. The protein alignments have been drawn to scale

**Fig 7.**
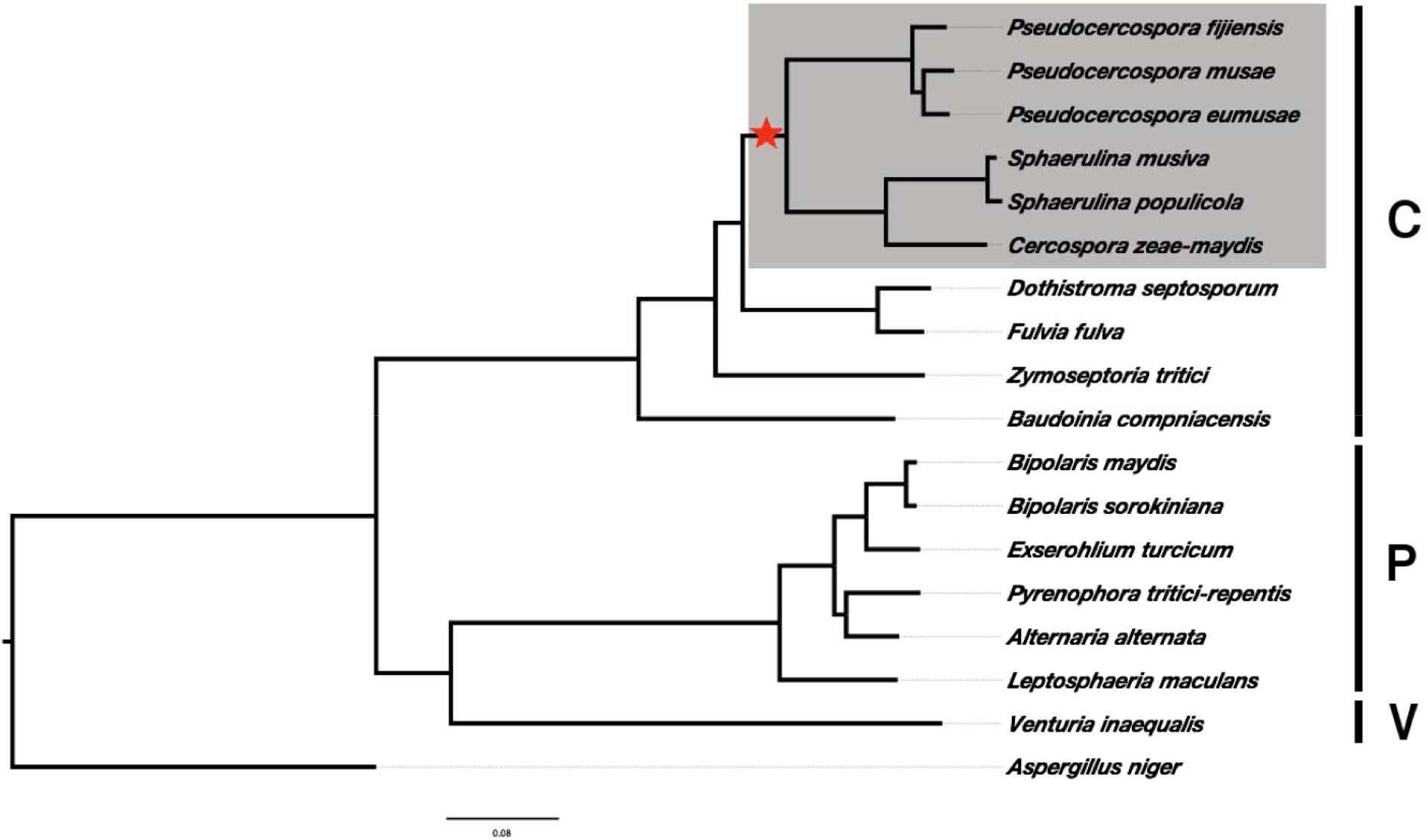
Presence of the *hAT*-mediated histone H3 amplification event in the family *Mycosphaerellaceae*, order Capnodiales of the fungal class Dothideomycetes. All six genomes available in the *Cercospora / Pseudocercospora / Sphaerulina* clade have multiple histone H3 copies that occur within *hAT* elements. The species highlighted in gray have multiple histone H3-like copies, whereas all other species lack the event. The star represents a divergence time estimate of 100 Mya (Rouxel *et al* 2011). Brackets to the right indicate the orders Capnodiales (C), Pleosporales (P) and Venturiales (V) in the class Dothideomycetes. A sequence of *Aspergillus niger* in the order Eurotiales of the class Eurotiomycetes was used as an outgroup

### The functional Histone H3 gene carries signatures of RIP

A consensus derived from ten H3-like sequences and its deRIP version was used to test whether RIP affected the original histone H3 coding sequence. These ten histone H3-like sequences spanned both the exon-exon junctions and covered at least 90% of the query sequence. The original H3 coding sequence had a GC content of 60%, whereas it ranged from 38–40% for the H3-like copies. The H3-like sequences were more similar to each other (89–98% identity) than they were to the original histone H3 coding sequence (51–55% identity).

A higher proportion of transition (Ti) mutations was seen in the H3-like sequences across a range of sites that were explored. The number of Ti mutations at variable as well as zero-, two-and four-fold degenerate sites was higher in the H3-like sequences as compared to the original histone H3 gene (Table 1). The direction of change for the Ti mutations (C>T, G>A) from the genomic histone H3 to repetitive H3-like sequence and *vice versa* was also evaluated. This analysis showed that the original histone H3 sequence also accumulated substitution mutations, even though the H3-like sequences had at least a 2x higher number of Ti mutations across all the classes of sites evaluated in both the deRIP and RIP consensus sequences (Table 1). As any substitution at a zero-fold degenerate site is non-synonymous, there were 13 (5.6%) sites that may have been changed in the original histone H3 sequence (Table 1) of *P. fijiensis*. However, the overall analysis showed that the histone H3 gene is under strong purifying selection with dN/dS ratio of 0.01429.

**Table 1.** Comparison of transition and transversion mutations among 513 total changes at 378 nucleotide positions among the aligned histone H3 sequences in the genome of *Pseudocercospora fijiensis*.

| Site category<sup>a</sup> | Mutation type<sup>b</sup> | Count (percent)<sup>c</sup> |  |
| --- | --- | --- | --- |
|  |  | deRIP consensus | RIP consensus |
| **Conserved** | —<sup>d</sup> | **186** | **180** |
| **Non-degenerate** |  | **234** | **233** |
| Conserved | — | 142 (60.7) | 135 (57.9) |
| Tv | Non-Synonymous | 56 (23.9) | 55 (23.6) |
| Ti from gH3 → rH3 | Non-Synonymous | 23 (9.8) | 32 (13.7) |
| Ti from rH3 → gH3 | Non-Synonymous | 13 (5.6) | 11 (4.7) |
| **Two-fold degenerate<sup>e</sup>** |  | **36** | **37** |
| Conserved | — | 20 (55.6) | 20 (54.1) |
| Tv | Non-Synonymous | 0 (0) | 1 (2.7) |
| Ti from gH3 → rH3 | Synonymous | 15 (41.7) | 15 (40.5) |
| Ti from rH3 → gH3 | Synonymous | 1 (2.8) | 1 (2.7) |
| **Four-fold degenerate** |  | **51** | **51** |
| Conserved | — | 8 (15.7) | 9 (17.6) |
| Tv | Synonymous | 21 (41.2) | 20 (39.2) |
| Ti from gH3 → rH3 | Synonymous | 16 (31.4) | 16 (31.4) |
| Ti from rH3 → gH3 | Synonymous | 6 (11.8) | 6 (11.8) |
| **Variable** |  | **192** | **198** |
| Tv | — | 100 (52.1) | 99 (50) |
| Ti from gH3 → rH3 | — | 70 (36.5) | 80 (40.4) |
| Ti from rH3 → gH3 | — | 22 (11.5) | 19 (9.6) |

<sup>a</sup> Non-degenerate sites are those at which any change in nucleotide results in an amino acid substitution; Two-fold degenerate sites are those at which only two of the four possible nucleotides specify the same amino acid; Four-fold degenerate sites are those at which all four nucleotides specify the same amino acid; Variable sites are those where the degeneracy status varies among the sequences being compared; Tv: Transversion mutations; Ti: Transition mutations; gH3: Original histone H3 sequence; rH3: H3-like sequences in transposable elements; The direction of change, either from gene (g) to repeat (r) or from repeat to gene is indicated by an arrow.
<sup>b</sup> Synonymous mutations cause no change in amino acid; Non-synonymous mutations lead to an amino acid substitution.
<sup>c</sup> Compared to a consensus sequence derived with repeat-induced point mutations present (RIP consensus) or after they were removed (deRIP consensus).
<sup>d</sup> Not applicable.
<sup>e</sup> Includes the only three-fold degenerate change in an isoleucine codon.

### Occurrence of histone H3 capture across the Dothideomycetes phylogeny

In addition to *P. fijiensis*, the histone H3 transduplication was identified in the genomes of five other species, viz., *Cercospora zeae-maydis, Pseudocercospora eumusae, P. musae, Sphaerulina musiva* and *S. populicola* among 12 Dothideomycetes genomes available in the family *Mycosphaerellaceae* (see Methods). Based on the phylogeny, it appears that a single histone H3 amplification event occurred prior to the split of the *Pseudocercospora*/*Cercospora*/*Septoria* clade from the other members of this family (Figure 8), which was estimated previously to have taken place approximately 100 Mya [23]. In addition to the 784 H3-like copies in *P. fijiensis*, a total of 242, 135, 520, 186 and 160 copies of H3-like sequences were identified in the repetitive fractions of the *C. zeae-maydis, P. eumusae, P. musae, S. musiva* and *S. populicola* genomes, respectively. As with *P. fijiensis*, a clear CA↔TA dinucleotide bias was observed in the repetitive elements carrying the histone H3-like sequences among these other fungal genomes. All of the extra copies contained premature stop codons due to RIP that would inactivate their function, except for a single, presumed original which contained an intact reading frame. Due to the fragmented nature of repeats that carry the histone H3-like sequences, the *hAT* domain could be identified only in *C. zeae-maydis, P. eumusae* and *S. musiva*, whereas TIR and TSD sequences could only be identified in the *C. zeae-maydis* genome.

Although a stretch of up to ten genes adjacent to the original histone H3 gene was syntenic between the *P. fijiensis* genome and those of the other five species, no synteny was found in the genomic regions around any of the histone H3-like copies. As expected, a stretch of nine or ten genes (including the original histone H3) was syntenic and collinear between *P. fijiensis* and the two closely related banana pathogens, *P. eumusae*, and *P. musae*, respectively, whereas the *Cercospora* and *Sphaerulina* genomes had eight genes that were in mesosynteny [24] with the *P. fijiensis* genome.

## Discussion

Previous analyses have shown that amplification of genes or gene fragments can have huge effects on genome architecture and evolution. However, as far as we know this is the first analysis in which a housekeeping gene has been amplified to a high copy number as part of a transposable element, yet all of the copies were inactivated, except for the original. This phenomenon resulted in the genome evolving to a much larger size due to the accumulation of numerous zombie copies of inactivated gene fragments. Capture and amplification of a transcript of the housekeeping gene histone H3 as part of a *hAT* DNA (class II) transposon was identified in six of 12 genomes tested in the family *Mycosphaerellaceae*, order Capnodiales of the fungal class Dothideomycetes. In each species all copies were inactivated by RIP, except for the presumed original, leading to one active gene plus hundreds of zombie copies scattered throughout the genome. Acquisition of a partial or complete copy of a gene between the termini of a DNA transposon and its subsequent amplification in a genome is termed ‘transduplication’. A *hAT* TE-mediated transduplication of the histone H3 coding sequence has been documented previously in the wheat pathogen *Pyrenophora tritici-repentis* (Ptr) [21], another Dothideomycete in the order Pleosporales. The occurrence of multiple, putatively functional H3-like copies in the Ptr genome appears to be the result of a recent and independent event, as evident from the paucity of mutations in the repetitive sequences and the presence of 12 identical copies in its genome [21].

RIP adds another layer of complexity to the possible outcomes of transposon-mediated gene captures and amplifications in fungi. With RIP, the rate at which the function of amplified genes is lost depends on several factors, including the number of codons amenable to RIP damage, the frequency of sexual reproduction (because RIP only occurs during meiosis) and the efficacy of the RIP machinery [17], which can vary by species. Length and sequence identity are two additional factors that affect RIP efficiency. Length of the *P. fijiensis* histone H3 coding sequence (411 bp) was just above the minimum cutoff (∼400 bp) required for recognition by the RIP machinery. Moreover, being a part of the longer hAT element, the H3-like retrocopies were more prone to RIP damage. The original histone H3 gene could have avoided RIP damage as it did not have a contiguous match of 400 bp with the retrocopy. However, directional mutations were observed in the histone H3 gene, suggesting that the length requirements for recognition by the RIP machinery may vary by species. Additionally, purifying selection would have played a role in maintaining the function of the full-length histone H3 gene; any mutations affecting function of the histones would likely be lethal and, therefore, be rapidly eliminated.

In the absence of RIP, the redundant gene copies after every amplification are free to evolve under different, and possibly relaxed, constraints. Even though transduplications lack promoter elements, moved gene fragments potentially can be expressed if inserted near regulatory elements [25] to obtain new functions. In plants, both transcription and translation of gene fragments transduced by Pack-MULEs, another type of class II transposon, have been demonstrated [26]. Besides the expression of processed pseudogenes, lack of RIP coupled with occurrence of multiple identical transcripts could also lead to post-transcriptional regulation of the original gene [27].

With the high frequency and efficiency of RIP in *P. fijiensis*, it seems highly unlikely that any of the duplicated histone H3 copies will contribute to future gene function in this fungus. Instead, all of the copies appear to have become rapidly pseudogenized. Similarly, RIP-induced mutations in an avirulence gene from *Leptosphaeria maculans* have been linked to the breakdown of major-gene-mediated resistance in *Brassica napus* [28]. This contrasts with organisms that lack RIP, where duplicated sequences often contribute to the standing genetic variation. In addition to the *Alu* repeats in humans [3], many other instances of TE or gene amplifications are known in different animals and plants [29,30]. For example, novel transcripts and proteins generated by pack-MULEs in rice undergo purifying selection and are maintained in its genome [26].

Two hypotheses could explain the occurrence of a single, intact copy of the histone H3 gene in a sea of inactivated, partial zombie copies. The first is that the original gene was protected from RIP, either due to its position within the genome or possibly because the duplicated fragment was too small to trigger recognition of the original gene. However, because there did appear to be RIP-induced changes in the original gene it seems much more likely that it was subject to RIP along with the copies but that any defects affecting function were eliminated by selection. RIP introduces bi-directional changes in all repetitive sequences in a genome, i.e., while the duplicated TEs accumulate RIP-introduced mutations, similar changes also can occur in the original histone H3 gene. Following its amplification by *hAT* transposition, histone H3, a housekeeping gene that is under strong negative selection, would suddenly become subjected to numerous transition mutations. Repetitive sequences that have more than 80% similarity continue to be targeted by the RIP machinery during every meiotic cycle [31]. Several meiotic cycles would be required before all copies of the repetitive H3-like genes became sufficiently diverged to completely disengage the original histone H3 gene from RIP damage. During this time, the original histone H3 gene most likely was affected by RIP. However, the strong purifying selection exerted on this gene appears to have eliminated any change that could affect its function.

Transduplication of the histone H3 gene in the *Mycosphaerellaceae* appears to be a relatively old event that most likely occurred in a common ancestor prior to the split of the *Pseudocercospora* and *Cercospora/Sphaerulina* lineages about 100 Mya [23]. Subsequent to this divergence, the genomes of the *Pseudocercospora* clade may have experienced one or more repeat-mediated expansions, resulting in a near doubling of their genomes compared to the average sizes of those from other Ascomycetes. The increased repetitive contents in *P. fijiensis* and *P. musae* are mirrored by the highest copy numbers of H3-like sequences in these two genomes. The cause of the relaxation of genome defense mechanisms that drove TE expansion in this clade is not known. Long periods of asexual reproduction could allow transposons to escape RIP, but this seems unlikely in *P. fijiensis* where the sexual stage is an integral part of the life cycle. Differences in copy numbers also may be affected slightly by the sequencing platform and the downstream assembly algorithms, leading to many poorly assembled copies of repetitive elements. However, this seems unlikely as the highest copy number of H3-like sequences (784) is found in the most well assembled genome, *P. fijiensis* (56 scaffolds; N50: 6Mb), i.e., copy number is not a proxy for poorly assembled genomes. If there is a bias, it would be in under-reporting of histone H3-like copies in genomes assembled from short sequencing reads.

Within the class Dothideomycetes there now exist three examples of independent TE-mediated amplifications that resulted in different outcomes for gene function and genome evolution. In the wheat pathogen *Z. tritici* in the order Capnodiales, a single-copy DNA methyltransferase gene was amplified to more than 20 copies, most likely through capture of a transcript followed by exchange among telomeres [19]. All copies were inactivated by RIP, including the original, leading to a loss of cytosine methylation in *Z. tritici*. This was postulated to be a recent event, as cytosine methylation was detected in very close relatives from wild grass hosts that are thought to have diverged within the past 10,000 years.

The second event was transduplication of the histone H3 gene in Ptr, in the order Pleosporales. This event also appeared to be very recent and involved capture of a transcript, but the amplification to tens of copies occurred through duplication and movement of a *hAT* transposon [21]. Here, RIP appears to be very inefficient or lacking, leaving multiple, potentially functional copies of the histone H3 gene.

The third event, initially identified in *P. fijiensis*, also involved capture of a histone H3 transcript by a *hAT* transposon, but unlike the other two examples appears to be very ancient, having originated in a common ancestor before the divergence of the *Cercospora/Pseudocercospora/Sphaerulina* clade from other species in the order Capnodiales. In this case, all of the copies have been heavily mutated and inactivated by RIP, except for the original, leading to a single functional copy of the histone H3 gene, and leaving repetitive regions that are graveyards of pseudogenized histone H3-like sequences. The original copy also was affected by RIP, but not to the point of altering the reading frame in a way that would prevent function. This most likely reflects the action of purifying selection to maintain the essential function of the histone H3 protein.

The three cases of gene amplification in the Dothideomycetes had different effects on genome evolution. In *Pyrenophora tritici*-*repentis*, amplification of the histone H3 gene and low efficiency or fewer cycles of RIP led to many transcriptionally active copies [21]. In the case of *Z. tritici*, where the original copy of a DNA methyltransferase gene as well as the amplified copies were mutated by RIP, the protein is not required and the fungus clearly can survive without cytosine methylation; presumably other types of methylation can compensate for the loss of this function [19]. In *P. fijiensis*, retention of the original histone H3 gene could have occurred for two reasons: 1) it is essential and its function cannot be lost so that all sexual (i.e., post-RIP) progeny with the original histone H3 gene mutated to the point of inactivation will not survive; and 2) the part that overlapped with the gene was too small to be targeted by RIP, so it retained fewer changes. Thus, the co-existence of gene amplification events and targeting of the RIP machinery to repetitive elements could lead to very different and unpredictable outcomes that impact both the function and evolution of fungal genomes.

## Conclusions

Previous analyses of repetitive sequences in fungal genomes identified two cases where genes were amplified to many copies but had different outcomes. In the first, all copies of a DNA methyltransferase gene in the wheat pathogen *Zymoseptoria tritici* (synonym *Mycosphaerella graminicola*) including the original were inactivated by repeat-induced point mutation (RIP), a genome-defense mechanism specific to fungi, leading to a loss of cytosine methylation in that species. The second case involved a different wheat pathogen, *Pyrenophora tritici-repentis*, in which RIP effects are lower, where capture and amplification of a histone H3 gene in a *hAT* DNA transposon led to multiple putatively active copies. Here a third case is identified, in which parts of a histone H3 gene were amplified to hundreds of copies as part of a *hAT* transposon, but all of the copies were highly mutated and inactivated by RIP, except for the original, leading to a greatly expanded genome but no additional functional copies of the gene. In contrast to the first two examples, this third case appears to be relatively ancient, and function of the original gene most likely was retained by strong purifying selection in spite of evident damage from RIP. These results demonstrate the variable effects that gene amplifications can have on the structure and evolution of genomes. The final outcome depends on the interplay of multiple factors that cannot be predicted without a much better understanding of genome biology.

## Methods

### Identification of the histone H3 amplification event

During the characterization of the *P. fijiensis* repetitive fraction [8], a histone H3-like sequence was identified in six families of repeats. The consensus repeats from these families have overlapping ends, i.e., they represent one repeat family that could be merged into a single contig. The original histone H3 protein sequence was then used to search the *P. fijiensis* genome using tBLASTn [32] to determine copy number. A similar search was used to identify H3-like sequences in the repetitive elements, and copy number per element was determined.

### Annotation of repeat families carrying the H3-like sequences

All of the repeat elements in the six families were annotated using TransposonPSI [33]. Repeat elements were aligned using clustalw [34] and the alignment was curated manually before RIP analysis [18]. Overlapping repeat-element sequences were merged irrespective of family, as the family delineations by repeat-finding programs are arbitrary. A set of four criteria-element length, presence of *hAT* domain, Terminal Inverted Repeat (TIR) and Target Site Duplication (TSD) -was used to identify the full-length copies in the merged repeat-element dataset. RIP analysis was also repeated on the subset of full-length repeat elements, each of which contained two copies of a histone H3-like gene.

### Histone H3-like sequence analysis

The full-length dataset was used to determine whether the genomic or coding histone H3 sequence was captured. Both of the H3-like copies present in each element were examined. Initially, the high-scoring segment pairs (HSP) resulting from the default tBLASTn output between the RIPped repeat elements and the original histone H3 protein were analyzed for the presence of exon-exon junctions. The multiple sequence alignment was deRIPped using the deRIP module in RIPCAL, which scans the alignment for polymorphic sites containing transition mutations (C/T or G/A) and reverses the effect of RIP. Manual curation was necessary to revert stop codons in coding sequences, as some sites were completely RIPped. To determine the directionality of the RIP mutations, near full-length H3-like sequences present in the repeat elements were aligned to the complete, original histone H3 coding sequence using RevTrans v.1.4 [35] and this alignment was visualized using MEGA v.6.06 [36]. Additionally, histone HMM was used with HMMER [37] to check the RIPped, *in silico* deRIPped, and manually curated deRIPped repeat sequences.

### Amplification of histone H3-like sequences in other fungi

The *P. fijiensis* histone H3 protein sequence was used to search 12 fungal genomes in the family *Mycosphaerellaceae* using tBLASTn [32]. Ten genome sequences obtained from the JGI Fungal Genome Portal (http://genome.jgi.doe.gov/programs/fungi/index.jsf), *Cercospora zeae-maydis* (abbreviated as Czm), *Dothistroma septosporum* (Dse), *Passalora fulva* (Pfu), *Pseudocercospora fijiensis* (Pfi), *Sphaerulina musiva* (Smu), *S. populicola* (Spo), *Zasmidium cellare* (Zce), *Zymoseptoria ardabiliae* (Zar), *Z. pseudotritici* (Zps) and *Z. tritici* (Ztr), and two additional genomes obtained from the NCBI, *Pseudocercospora eumusae* (Peu; GenBank accession number GCA_001578235.1) and *P. musae* (Pmu; GCA_001578225.1), were scanned for the histone H3 amplification event. Protein sets for the above-mentioned genomes and nine additional genomes, those of *Baudoinia compniacensis* in the order Capnodiales, *Alternaria alternata, Bipolaris sorokiniana* [synonym: *Cochliobolus sativus*], *B. maydis* [*C. heterostrophus* isolate C5], *Leptosphaeria maculans, Pyrenophora tritici-repentis, Setosphaeria turcica*, in the order Pleosporales, and *Venturia inaequalis* in the order Venturiales, all of the class Dothideomycetes, plus *Aspergillus niger* in the order Eurotiales of the class Eurotiomycetes as an outgroup, also were downloaded for analysis.

The protein datasets from 18 genomes, except for Zar, Zce, and Zps were used for an all-versus-all BLAST. The BLAST output was analyzed using OrthoMCL [38] to identify 2,220 one-to-one orthologous clusters (OrthoMCL inflation value of I = 1.5). These orthologous cluster sequences were then aligned using ClustalX [34]. For each alignment, conserved blocks were identified using Gblocks [39] at default settings. ProtTest [40] was used subsequently to identify the best model of protein evolution for each alignment using the Akaike Information Criterion (AIC). The protein alignments were then concatenated and used for generating a maximum likelihood (ML) species phylogeny using RAxML [41]. A similar ML phylogeny using the original Histone H3 protein sequences from these 18 genomes was generated using RAxML and the dN/dS ratio was determined using PAML [42].

A stretch of 10 kb of the sequence up-and down-stream of the original *P. fijiensis* histone H3 gene was used to determine the extent of synteny with the five most closely related genomes (Czm, Peu, Pmu, Smu and Spo). Additionally, a search for orthologous repetitive elements carrying the H3-like sequences also was conducted.

## Acknowledgements

This work was funded in part by USDA-ARS CRIS project 5020-22000-017-00D. Genomic sequencing of *P. fijiensis* and several other species was performed at the U. S. Department of Energy’s Joint Genome Institute through the Community Sequencing Program (www.jgi.doe.gov/csp/) and are publicly available.

## Availability of data and materials

Genome sequences for the 19 fungi analyzed are available from the JGI Mycocosm database (http://genome.jgi.doe.gov/programs/fungi/index.jsf) and two genomes are available from GenBank (GCA_001578235.1; GCA_001578225.1).

## Authors’ contributions

BD co-designed the project, performed all of the analyses, made the figures, wrote the initial draft of the manuscript and edited subsequent versions. GHJK co-wrote the grant and provided the DNA that generated the genome sequence for *P. fijiensis*. RH wrote the grant and provided the materials that generated the sequences of the two *Sphaerulina* species. BHB provided funding and materials to sequence the genome of *C. zeae-maydis*. SBG co-wrote the grant to sequence the genome of *P. fijiensis*, co-designed the project, wrote portions and revised and edited the entire manuscript. All authors read and approved the final manuscript.

## Ethics approval and consent to participate

Not applicable.

## Consent for publication

Not applicable.

## Additional files

**Additional file 1: Figure S1.** Analysis of di-nucleotide bias in the hisotne H3-carrying *hAT* DNA transposon family in *Pseudocercospora fijiensis*. Alignment of 917 repeat sequences showed a clear CA<->TA RIP bias

**Additional file 2: Figure S2.** Termini from the full-length *hAT* element repeats that carry histone H3-like sequences in *Cercospora zeae-maydis*. This alignment highlights 18-bp terminal inverted repeat (TIR) and 8-bp target site duplication (TSD) sequences. TIR and TSD are separated by a single gap (−) symbol

**Additional file 3: Figure S3.** Organization of the histone H3 gene region in six Dothideomycete genomes. A 10-kb sequence up-and down-stream of the histone H3 gene in the Pfi genome was used to do a pair-wise comparison with five other genomes: Pmu, Peu, Czm, Smu and Spo. The 20-kb region from Pfi is syntenic with Pmu and Peu, whereas genes from this region are dispersed across the scaffolds in Czm, Smu and Spo genomes (mesosynteny as defined by [24]). Pfi sequence was used as reference and shown on the X axis. Protein similarity between the two genomes is shown by the colored lines in the plots

## References

1. Munoz-Lopez M, Garcia-Perez JL. DNA transposons: nature and applications in genomics. Curr. Genomics. Netherlands; 2010;11:115–28.

2. Kazazian HHJ. Mobile elements: drivers of genome evolution. Science. United States; 2004;303:1626–32.

3. Deininger P. Alu elements: know the SINEs. Genome Biol. [Internet]. 2011;12:236. Available from: http://dx.doi.org/10.1186/gb-2011-12-12-236

4. Doebley J, Stec A, Gustus C. teosinte branched1 and the origin of maize: evidence for epistasis and the evolution of dominance. Genetics. United States; 1995;141:333–46.

5. Studer A, Zhao Q, Ross-Ibarra J, Doebley J. Identification of a functional transposon insertion in the maize domestication gene tb1. Nat. Genet. United States; 2011;43:1160–3.

6. Xiao H, Jiang N, Schaffner E, Stockinger EJ, van der Knaap E. A Retrotransposon-Mediated Gene Duplication Underlies Morphological Variation of Tomato Fruit. Science (80-.). [Internet]. 2008;319:1527–30. Available from: http://www.sciencemag.org/cgi/doi/10.1126/science.1153040

7. Spanu PD, Abbott JC, Amselem J, Burgis TA, Soanes DM, Stuber K, et al. Genome expansion and gene loss in powdery mildew fungi reveal tradeoffs in extreme parasitism. Science. United States; 2010;330:1543–6.

8. Arango Isaza RE, Diaz-Trujillo C, Dhillon B, Aerts A, Carlier J, Crane CF, et al. Combating a global threat to a clonal crop: Banana Black Sigatoka pathogen Pseudocercospora fijiensis (Synonym Mycosphaerella fijiensis) genomes reveal clues for disease control. PLOS Genet. [Internet]. Public Library of Science; 2016;12:e1005876. Available from: https://doi.org/10.1371/journal.pgen.1005876

9. Grandaubert J, Lowe RGT, Soyer JL, Schoch CL, Van de Wouw AP, Fudal I, et al. Transposable element-assisted evolution and adaptation to host plant within the Leptosphaeria maculans-Leptosphaeria biglobosa species complex of fungal pathogens. BMC Genomics. England; 2014;15:891.

10. Sacristán S, Vigouroux M, Pedersen C, Skamnioti P, Thordal-Christensen H, Micali C, et al. Coevolution between a family of parasite virulence effectors and a class of LINE-1 retrotransposons. PLoS One [Internet]. Public Library of Science; 2009;4:e7463. Available from: https://doi.org/10.1371/journal.pone.0007463

11. Amselem J, Vigouroux M, Oberhaensli S, Brown JKM, Bindschedler L V, Skamnioti P, et al. Evolution of the EKA family of powdery mildew avirulence-effector genes from the ORF 1 of a LINE retrotransposon. BMC Genomics. England; 2015;16:917.

12. Orbach MJ, Farrall L, Sweigard JA, Chumley FG, Valent B. A telomeric avirulence gene determines efficacy for the rice blast resistance gene Pi-ta. Plant Cell. United States; 2000;12:2019–32.

13. Ma L-J, van der Does HC, Borkovich KA, Coleman JJ, Daboussi M-J, Di Pietro A, et al. Comparative genomics reveals mobile pathogenicity chromosomes in Fusarium. Nature [Internet]. Macmillan Publishers Limited. All rights reserved; 2010;464:367–73. Available from: http://dx.doi.org/10.1038/nature08850

14. Yoshida K, Saunders DGO, Mitsuoka C, Natsume S, Kosugi S, Saitoh H, et al. Host specialization of the blast fungus Magnaporthe oryzae is associated with dynamic gain and loss of genes linked to transposable elements. BMC Genomics [Internet]. 2016;17:370. Available from: http://dx.doi.org/10.1186/s12864-016-2690-6

15. Ku H-Y, Lin H. PIWI proteins and their interactors in piRNA biogenesis, germline development and gene expression. Natl. Sci. Rev. [Internet]. 2014;1:205–18. Available from: http://dx.doi.org/10.1093/nsr/nwu014

16. Selker EU, Cambareri EB, Jensen BC, Haack KR. Rearrangement of duplicated DNA in specialized cells of Neurospora. Cell. United States; 1987;51:741–52.

17. Galagan JE, Selker EU. RIP: the evolutionary cost of genome defense. Trends Genet. England; 2004;20:417–23.

18. Hane JK, Oliver RP. RIPCAL: a tool for alignment-based analysis of repeat-induced point mutations in fungal genomic sequences. BMC Bioinformatics [Internet]. 2008;9:478. Available from: http://dx.doi.org/10.1186/1471-2105-9-478

19. Dhillon B, Cavaletto JR, Wood K V, Goodwin SB. Accidental amplification and inactivation of a methyltransferase gene eliminates cytosine methylation in Mycosphaerella graminicola. Genetics [Internet]. 2010;186:67–77. Available from: http://www.genetics.org/content/186/1/67.abstract

20. Stukenbrock EH, Banke S, Javan-Nikkhah M, McDonald BA. Origin and domestication of the fungal wheat pathogen Mycosphaerella graminicola via sympatric speciation. Mol. Biol. Evol. United States; 2007;24:398–411.

21. Manning VA, Pandelova I, Dhillon B, Wilhelm LJ, Goodwin SB, Berlin AM, et al. Comparative genomics of a plant-pathogenic fungus, Pyrenophora tritici-repentis, reveals transduplication and the impact of repeat elements on pathogenicity and population divergence. G3 (Bethesda). United States; 2013;3:41–63.

22. Rubin E, Lithwick G, Levy AA. Structure and evolution of the hAT transposon superfamily. Genetics. United States; 2001;158:949–57.

23. Rouxel T, Grandaubert J, Hane JK, Hoede C, van de Wouw AP, Couloux A, et al. Effector diversification within compartments of the Leptosphaeria maculans genome affected by Repeat-Induced Point mutations. Nat. Commun. [Internet]. The Author(s); 2011;2:202. Available from: http://dx.doi.org/10.1038/ncomms1189

24. Hane JK, Rouxel T, Howlett BJ, Kema GHJ, Goodwin SB, Oliver RP. A novel mode of chromosomal evolution peculiar to filamentous Ascomycete fungi. Genome Biol. England; 2011;12:R45.

25. Sakai H, Koyanagi KO, Imanishi T, Itoh T, Gojobori T. Frequent emergence and functional resurrection of processed pseudogenes in the human and mouse genomes. Gene [Internet]. 2007;389:196–203. Available from: http://www.sciencedirect.com/science/article/pii/S0378111906007098

26. Hanada K, Vallejo V, Nobuta K, Slotkin RK, Lisch D, Meyers BC, et al. The functional role of Pack-MULEs in rice inferred from purifying selection and expression profile. Plant Cell [Internet]. American Society of Plant Biologists (ASPB); 2009;21:25–38. Available from: http://www.jstor.org/stable/40537459

27. Chiefari E, Iiritano S, Paonessa F, Le Pera I, Arcidiacono B, Filocamo M, et al. Pseudogene-mediated posttranscriptional silencing of HMGA1 can result in insulin resistance and type 2 diabetes. Nat. Commun. [Internet]. Nature Publishing Group, a division of Macmillan Publishers Limited. All Rights Reserved.; 2010;1:40. Available from: http://dx.doi.org/10.1038/ncomms1040

28. Fudal I, Ross S, Brun H, Besnard A-L, Ermel M, Kuhn M-L, et al. Repeat-induced point mutation (RIP) as an alternative mechanism of evolution toward virulence in Leptosphaeria maculans. Mol. Plant. Microbe. Interact. United States; 2009;22:932–41.

29. Lisch D. How important are transposons for plant evolution? Nat Rev Genet [Internet]. Nature Publishing Group, a division of Macmillan Publishers Limited. All Rights Reserved.; 2013;14:49–61. Available from: http://dx.doi.org/10.1038/nrg3374

30. Warren IA, Naville M, Chalopin D, Levin P, Berger CS, Galiana D, et al. Evolutionary impact of transposable elements on genomic diversity and lineage-specific innovation in vertebrates. Chromosom. Res. Netherlands; 2015;23:505–31.

31. Cambareri E, Singer M, Selker E. Recurrence of repeat-induced point mutation (RIP) in Neurospora crassa. Genetics. 1991;127:699–710.

32. Altschul SF, Gish W, Miller W, Myers EW, Lipman DJ. Basic local alignment search tool. J. Mol. Biol. England; 1990;215:403–10.

33. Hass B. TransposonPSI: An application of PSI-Blast to mine (retro−)transposon ORF homologies. [Internet]. 2010. Available from: http://transposonpsi.sourceforge.net/

34. Larkin MA, Blackshields G, Brown NP, Chenna R, McGettigan PA, McWilliam H, et al. Clustal W and Clustal X version 2.0. Bioinformatics [Internet]. 2007;23:2947–8. Available from: http://dx.doi.org/10.1093/bioinformatics/btm404

35. Wernersson R, Pedersen AG. RevTrans: Multiple alignment of coding DNA from aligned amino acid sequences. Nucleic Acids Res. England; 2003;31:3537–9.

36. Tamura K, Stecher G, Peterson D, Filipski A, Kumar S. MEGA6: Molecular Evolutionary Genetics Analysis Version 6.0. Mol. Biol. Evol. [Internet]. Oxford University Press; 2013;30:2725–9. Available from: http://www.ncbi.nlm.nih.gov/pmc/articles/PMC3840312/

37. Johnson LS, Eddy SR, Portugaly E. Hidden Markov model speed heuristic and iterative HMM search procedure. BMC Bioinformatics [Internet]. 2010;11:431. Available from: http://dx.doi.org/10.1186/1471-2105-11-431

38. Li L, Stoeckert CJ, Roos DS. OrthoMCL: identification of ortholog groups for eukaryotic genomes. Genome Res [Internet]. 2003;13:2178–89. Available from: http://dx.doi.org/10.1101/gr.1224503

39. Castresana J. Selection of conserved blocks from multiple alignments for their use in phylogenetic analysis. Mol. Biol. Evol. United States; 2000;17:540–52.

40. Abascal F, Zardoya R, Posada D. ProtTest: selection of best-fit models of protein evolution. Bioinformatics [Internet]. 2005;21:2104–5. Available from: http://dx.doi.org/10.1093/bioinformatics/bti263

41. Stamatakis A. RAxML version 8: a tool for phylogenetic analysis and post-analysis of large phylogenies. Bioinformatics [Internet]. Oxford University Press; 2014;30:1312–3. Available from: http://www.ncbi.nlm.nih.gov/pmc/articles/PMC3998144/

42. Yang Z. PAML: a program package for phylogenetic analysis by maximum likelihood. Comput. Appl. Biosci. England; 1997;13:555–6.

