## Supplementary figures and images for "Variable Genome Evolution in Fungi After Transposon-Mediated Amplification of a Housekeeping Gene"

### Supplemental Figure 1

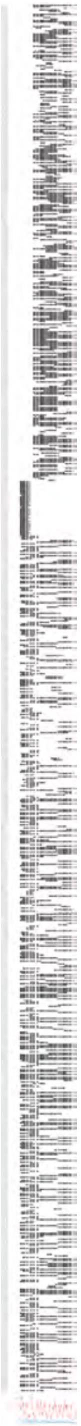

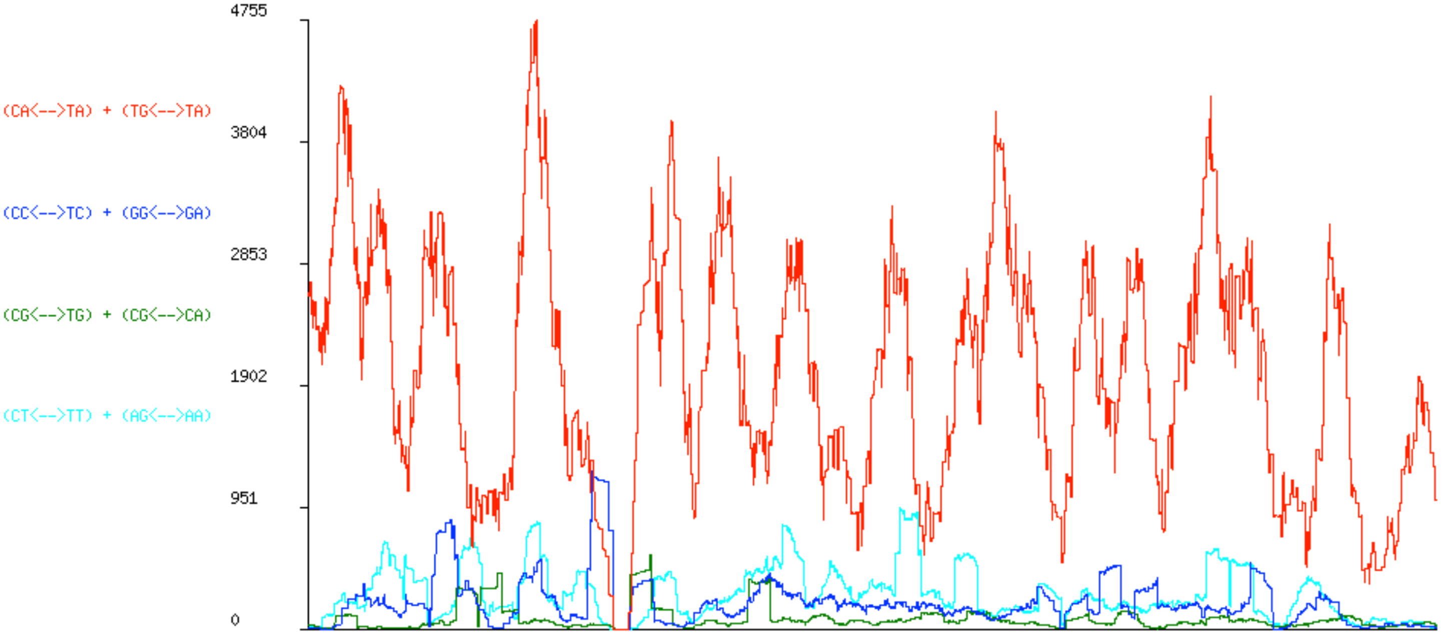

### Supplemental Figure 2

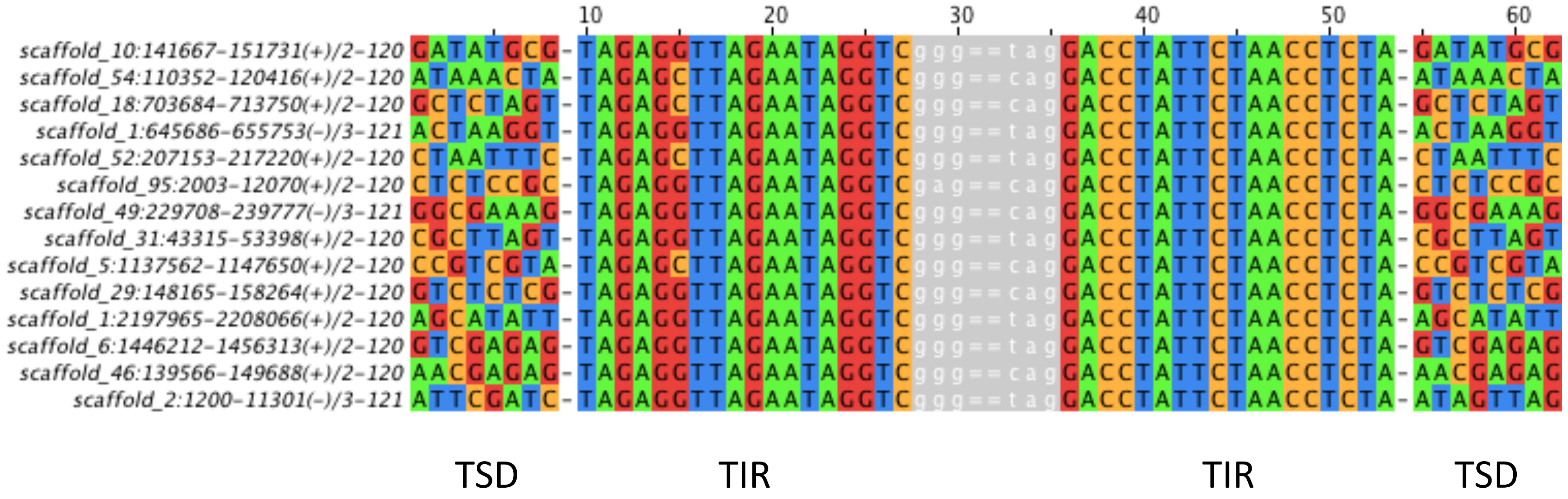

### Supplemental Figure 3

S3

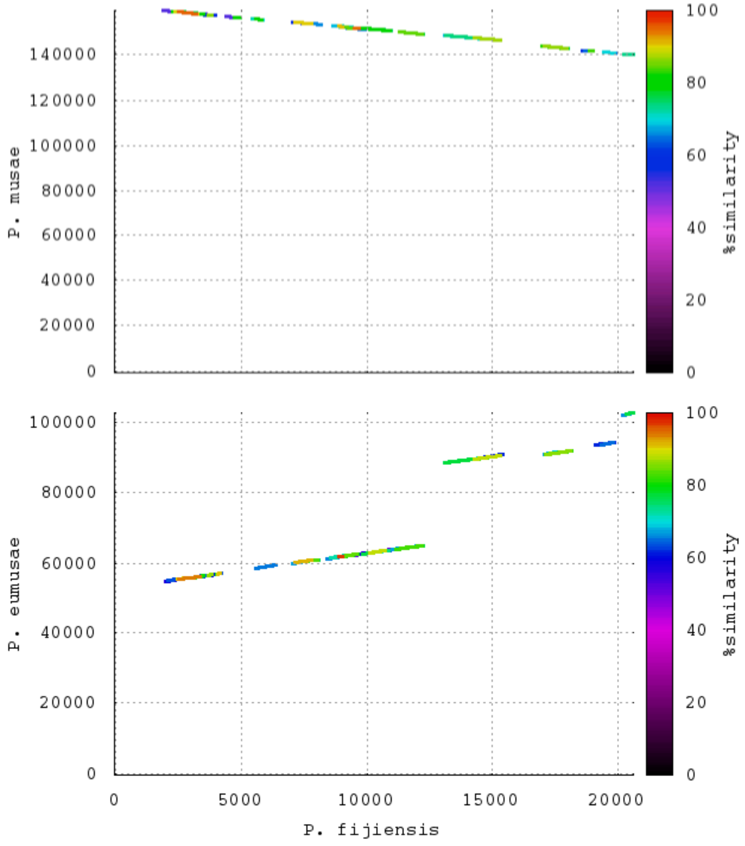


Synteny in Pfi/Pmu; Pfi/Peu

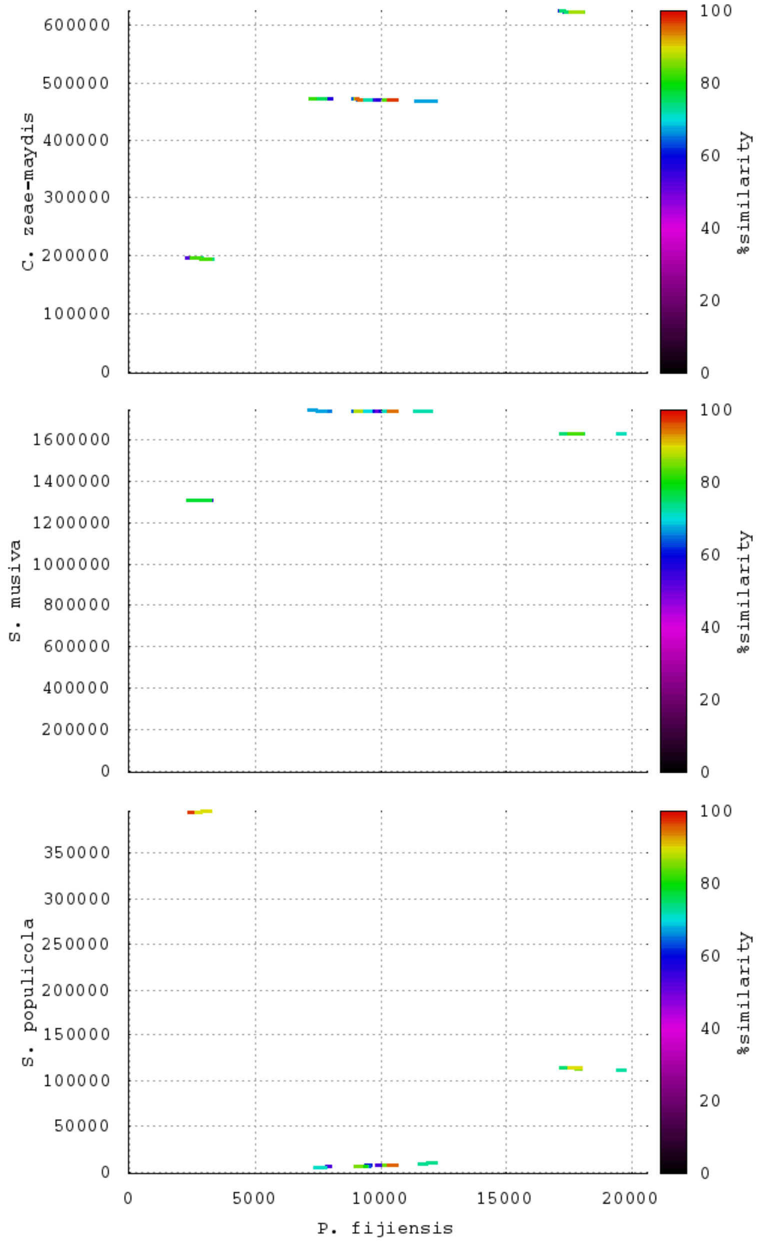


Mesosynteny in Pfi/Czm; Pfi/Smu; Pfi/Spo
